# BNT162b2-Elicited Neutralization of Delta Plus, Lambda, and Other Variants

**DOI:** 10.1101/2021.09.13.460163

**Authors:** Jianying Liu, Yang Liu, Hongjie Xia, Jing Zou, Scott C. Weaver, Kena A. Swanson, Hui Cai, Mark Cutler, David Cooper, Alexander Muik, Kathrin U. Jansen, Ugur Sahin, Xuping Xie, Philip R. Dormitzer, Pei-Yong Shi

## Abstract

BNT162b2-elicited human sera are known to neutralize the currently dominant Delta SARS-CoV-2 variant. Here, we report the ability of 20 human sera, drawn 2 or 4 weeks after two doses of BNT162b2, to neutralize USA-WA1/2020 SARS-CoV-2 bearing variant spikes from Delta plus (Delta-AY.1, Delta-AY.2), Delta-Δ144 (Delta with the Y144 deletion of the Alpha variant), Lambda, and B. 1.1.519 lineage viruses. Geometric mean plaque reduction neutralization titers against Delta-AY.1, Delta-AY.2, and Delta-Δ144 viruses are slightly lower than against USA-WA1/2020, but all sera neutralize the variant viruses to titers of ≥80.

Neutralization titers against Lambda and B. 1.1.519 variants and against USA-WA1/2020 are equivalent. The susceptibility of Delta plus, Lambda, and other variants to neutralization by the sera indicates that antigenic change has not led to virus escape from vaccine-elicited neutralizing antibodies and supports ongoing mass immunization with BNT162b2 to control the variants and to minimize the emergence of new variants.

## Introduction

As of August 13, 2021, severe acute respiratory syndrome coronavirus 2 (SARS-CoV-2) has caused over 205 million infections and more than 4.3 million deaths due to coronavirus disease 2019 (COVID-19; https://coronavirus.jhu.edu/). Since its emergence in late 2019, SARS-CoV-2 has accumulated mutations, leading to variants with higher transmission, more efficient replication, and potentially immune evasion.^1–5^ Many of these mutations have occurred in the viral spike glycoprotein, which is responsible for binding to the host receptor, angiotensin-converting enzyme 2 (ACE2), during virus entry. Based on the effects of mutations on viral transmission, disease severity, and clinical diagnosis, the World Health Organization (WHO) has categorized SARS-CoV-2 strains into “variants of concern (VOC),” “variants of interest (VOI),” and “variants of alert (VOA)” (https://www.who.int/en/activities/tracking-SARS-CoV-2-variants/). Currently, VOC include Alpha (B.1.1.7), Beta (B. 1.351), Gamma (P.1), and Delta (B.1.617.2); VOI include Eta (B.1.525), Iota (B.1.126), Kappa (B.1.617.1), and Lambda (C.37). As the pandemic continues, it is critical to monitor closely the new variants for their transmission, pathogenesis, and potential escape from vaccines and therapeutics.

BNT162b2 is an mRNA vaccine expressing the full-length prefusion spike glycoprotein of SARS-CoV-2, stabilized in the prefusion conformation.^6^ BNT162b2 has recently been approved for vaccination of individuals 16 years of age and older and has been authorized under emergency use provisions for immunization of those 12-15 years old by the US Food and Drug Administration. Although BNT162b2 mRNA encodes the original spike protein from the Wuhan isolate,^7^ the sera of those immunized with BNT162b2 can neutralize all tested variants, including the currently circulating Delta variant.^2,8–13^ However, some variants are less efficiently neutralized than others, with the Beta and Kappa variants showing the greatest decrease to date.^9,10,14^ The explosive recent spread of the Delta variant to 119 countries and its association with breakthrough infections in vaccinated people prompted us to examine the closely related Delta plus variants, such as (i) Delta-AY.1 (first detected in India and spread to 32 countries, including the USA); (ii) Delta-AY.2 (first detected in the USA and spread to 7 countries); and (iii) Delta-Δ144 (first detected in Vietnam and spread to 17 countries) (https://www.gisaid.org/hcov19-variants/). In addition to the Delta variants, the Lambda variant (C.37; first detected in Peru) has spread to 46 countries with high prevalence in South America; the B.1.1.519 variant has emerged and became dominant in Mexico during the first months of 2021 (https://www.gisaid.org/hcov19-variants/). Consequently, the WHO has designated Delta-AY.1 and Delta-AY.2 as VOC, Lambda as VOI, and B.1.1.519 as VOA. Here, we report BNT162b2 vaccine-elicited neutralization against these new variants.

## Results

We aimed to study the impact of antigenic variation in the SARS-CoV-2 spike glycoprotein on neutralization by antibodies elicited by the wild type (WT) spike glycoprotein encoded by BNT162b2 RNA. Therefore, we used a reverse genetic system to generate a panel of SARS-CoV-2 with a USA-WA1/2020 genetic background (a viral strain isolated in January 2020 and defined as WT) and spike glycoproteins from the newly emerged variants (**Extended data Fig. 1a**). Five chimeric SARS-CoV-2’s were prepared: (i) Delta-AY.1-spike with T19R, T95I, G142D, E156G, F157/R158 deletion (Δ157/158), W258L, K417N, L452R, T478K, K558N, D614G, P681R, and D950N mutations (GISAID accession ID: EPI_ISL_2676768); (ii) Delta-AY.2-spike with T19R, V70F, G142D, E156G, Δ157/158, A222V, K417N, L452R, T478K, D614G, P681R, D950N, and V1228L (GISAID accession ID: EPI_ISL_2527809); (iii) Delta-Δ144-spike (a Delta variant that has acquired a Y144-deletion from the Alpha variant) with T19R, G142D, Δ144, E156G, Δ157/158, A222V, L452R, T478K, D614G, P681R, and D950N (GISAID accession ID: EPI_ISL_2373110); (iv) Lambda-spike with G75V, T76I, R246-G252 deletion (Δ246-252), D253N, L452Q, F490S, D614G, and T859N (GISAID accession ID: EPI_ISL_1138413); and (v) B.1.1.519-spike with T478K, D614G, P681H, and T732A (GISAID accession ID: EPI_ISL_876555). The WT and all five chimeric viruses were rescued from infectious cDNA clones and titered by plaque assay on Vero E6 cells. All these viruses generated similar plaque morphologies (**Extended Data Fig. 1b**) and infectious titers of greater than 10^7^ plaque-forming units (PFU)/ml. The ratio of viral RNA to PFU was quantified for each virus; no significant differences in viral RNA-to-PFU ratios were detected between the WT and chimeric variant viruses (**Extended Data Fig. 1c**), indicating similar specific infectivities. Sequencing of viral stocks confirmed that there were no undesired mutations in the spike gene.

We compared the neutralization susceptibility of the chimeric variant SARS-CoV-2s to a panel of 20 sera collected from BTN162b2-immunized human participants in the pivotal clinical trial.^6,15^ As reported previously, the serum specimens were drawn 2 or 4 weeks after two immunizations with 30 μg of BNT162b2, spaced three weeks apart.^9,10^ Each serum was tested simultaneously for its 50% plaque reduction neutralizing titers (PRNT_50_) against the WT and chimeric variant viruses (**Extended Data Table 1**). All the sera neutralized the WT and all the mutant viruses with titers of 1:80 or higher (**Fig. 1**). The geometric mean neutralizing titers against the WT, Delta-AY.1-spike, Delta-AY.2-spike, Delta-Δ144-spike, Lambda-spike, and B.1.1.519-spike viruses were 520, 355, 394, 453, 597, and 640, respectively (**Fig. 1**). The results indicate that neutralization of Delta plus variants (Delta-AY.1, Delta-AY.2, and Delta-Δ144) is only modestly reduced relative to neutralization of WT virus. We previously reported a similar neutralization result for another Delta plus strain, Delta-AY.3 (B.1.617.2.v2),^8^ which currently accounts for 13.4% of clinical SARS-CoV-2 isolates in the US (https://covid.cdc.gov/covid-data-tracker/#variant-proportions). Neutralization of the Lambda and B.1.1.519 variants is not reduced relative to neutralization of WT virus. Thus, BNT162b2 immune sera efficiently neutralized all tested viruses.

**Figure 1.**
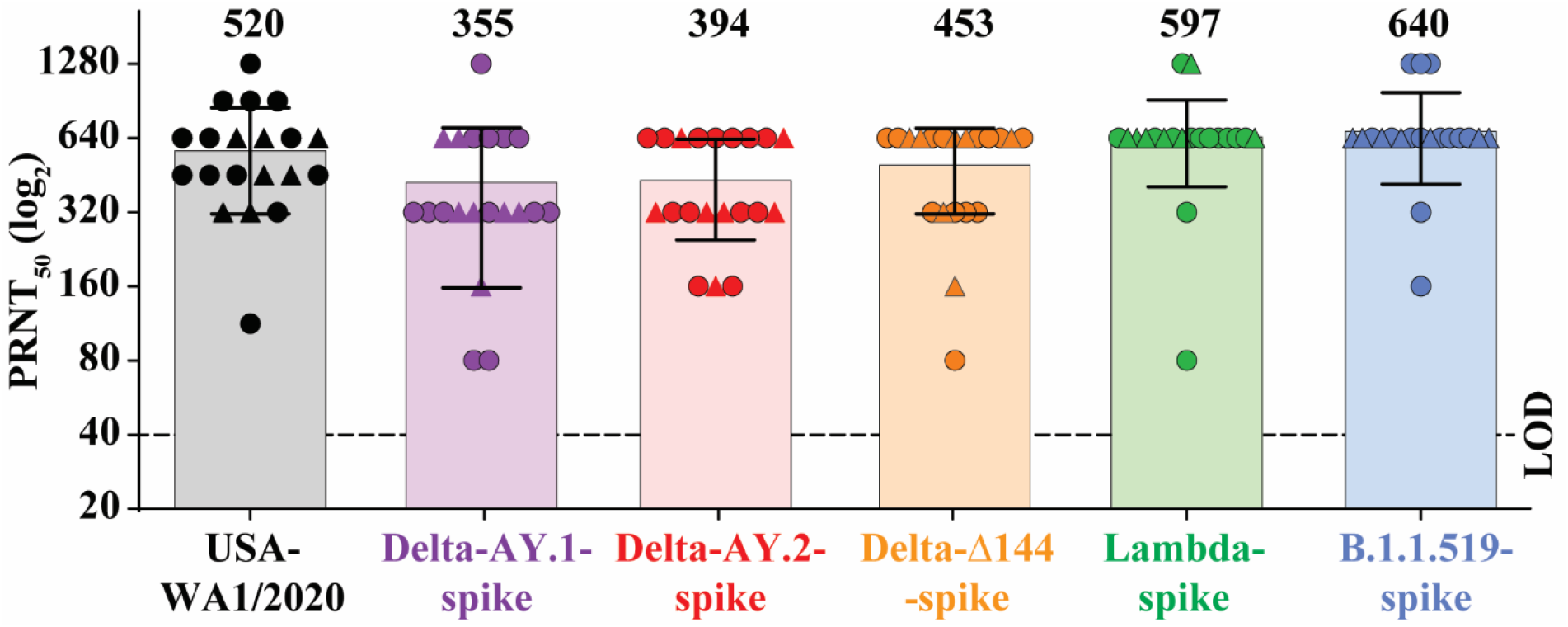
BNT162b2 vaccine-elicited neutralization of SARS-CoV-2 variants. The plot presents the PRNT_50_ titers of 20 human sera (drawn 2 or 4 weeks after two 30-μg doses of BNT162b2, spaced three weeks apart) against USA-WA1/2020 isolate and its chimeric viruses bearing distinct variant spikes. Serum samples obtained at 2 weeks or 4 weeks are represented by circles and triangles, respectively. Individual PRNT_50_ values are presented in **Extended Data Table 1**. Each data point represents the geometric mean PRNT_50_ against the indicated virus obtained with a serum specimen. The PRNT_50_ values were determined in duplicate assays, and the geometric means were calculated (n=20, pooled from two independent experiments). The bars and the numbers above the bars indicate geometric mean titers. The horizontal bars indicate 95% confidence intervals. The limit of detection (LOD) of the PRNT assay is 1:40 and indicated by a dashed line. Statistical analysis was performed using the two-tailed Wilcoxon matched-pairs signed-rank test. The statistical significances of the differences between geometric mean titers in the USA-WA1/2020 neutralization assay and in each variant virus neutralization assay with the same serum samples are as follows: *P* = 0.0264 for Delta-AY.1-spike; *P* < 0.030 for Delta-AY.2-spike; *P* = 0.255 for Delta-Δ144-spike; *P* = 0.193 for Lambdaspike and *P* = 0.007 for B.1.1.519-spike.

## Discussion

We have taken a systematic approach to measuring BNT162b2-elicted neutralization of newly emerged variants. We use a reverse genetic system to generate chimeric SARS-CoV-2’s bearing spikes from distinct variants through site-directed mutagenesis or DNA synthesis. This approach has two major advantages: (i) it allows us to examine new variants as soon as their sequences become available, and (ii) it measures the impact of variant spikes on neutralizing activity without being affected by mutations outside the spike gene. Non-spike mutations are not directly relevant to the selection of vaccine spike sequences. We test all chimeric variants for neutralization by the same panel of 20 sera from BNT162b2-vaccinated trial participants, enabling us to make well controlled comparisons longitudinally to inform vaccine decision making^8–10,16^.

Our previous and current results suggest that BNT162b2-vaccinated sera neutralize the Delta and Delta plus variants more efficiently than they neutralize the Beta variant. Real-world effectiveness of BNT162b2 against Beta variant-associated severe or fatal disease in Qatar and vaccine efficacy against Beta variant-associated COVID-19 in South Africa were both reported to have point estimates of 100%.^12,17^ These results indicate that the observed breakthrough infections of the Delta variant are not due to immune escape. Recent studies using primary human airway cultures and a human lung epithelial cell line suggest that the Delta variant has improved replication fitness through mutation P681R-enhanced protease cleavage of the fulllength spike to S1 and S2 subunits, as one mechanism leading to increased viral infection.^18,19^ Viral RNA loads in the oropharynx from Delta variant-infected patients were over 1,000-fold higher than those from the original Wuhan virus-infected individuals.^20^ Collectively, the results suggest that improved viral fitness due to more efficient furin cleavage, rather than immune escape, may account for breakthrough infections of the Delta variant in vaccinated people. More efficient furin cleavage of the influenza fusion protein, hemagglutinin, increases the viral pathogenicity to an even greater extent.^21^

One limitation of this study is the potential for mutations to alter neutralization by affecting spike function rather than antigenicity (*e.g*., mutation P681R that improves viral replication through enhanced spike processing), despite the variant viruses exhibiting specific infectivities similar to that of the original USA-WA1/2020 virus on Vero E6 cells. Another limitation is that the study focuses only on the effect of spike glycoprotein mutations on neutralization in cell culture. Mutations outside the spike gene may also alter viral replication and/or host immune response.

The susceptibility of Delta, Delta plus, Lambda, and other variants to BNT162b2-elicited neutralization indicates that antigenic change does not yet appear to be the major mechanism of increased Delta variant pathogenicity or spread. This finding suggests that changing the strain of the spike glycoprotein encoded by the vaccine may not be the most effective response to the emergence and spread of the Delta variant. Nevertheless, Pfizer and BioNTech are preparing for the possibility that a strain change may someday be necessary by testing prototype, antigenically updated vaccines. The data in this paper support the ongoing BNT162b2 mass immunization strategy to control the variants and to minimize the emergence of new variants. Increasing the vaccination rate in the population, together with implementing public health measures, remains the primary means to end the COVID-19 pandemic.

## Methods

### Cells

Vero E6 cells, an African green monkey kidney epithelial cell line (ATCC, Manassas, VA, USA), were cultured in Dulbecco’s modified Eagle’s medium (DMEM; Gibco/Thermo Fisher, Waltham, MA, USA) with 10% fetal bovine serum (FBS; HyClone Laboratories, South Logan, UT) plus 1% ampicillin/streptomycin (Gibco). The authenticity of Vero E6 cells was verified through STR profiling by ATCC. The cells were tested negative for mycoplasma.

### Construction of chimeric SARS-CoV-2s with variant spikes

All spike mutations from variants were engineered into infectious cDNA clones of an early SARS-CoV-2 isolate, USA-WA1/2020, using a standard PCR-based mutagenesis method.^22^ The protocol for construction of recombinant SARS-CoV-2 was reported previously.^23^ The full-length cDNAs of viral genomes containing the variant spike mutations were assembled by *in vitro* ligation. The resulting genome-length cDNAs served as templates for *in vitro* transcription of full-length viral RNAs. The full-length viral RNA transcripts were electroporated into Vero E6 cells. On day 2 post electroporation (when the electroporated cells developed cytopathic effects due to recombinant virus production and replication), the original viral stocks (P0) were harvested from culture medium. The P0 viruses were amplified on Vero E6 cells for another round to produce working viral stocks (P1). The infectious titers of the P1 viruses were quantified by plaque assay on Vero E6 cells.^22^ The complete spike genes from the P1 viruses were sequenced to ensure no undesired mutations. The P1 viruses were used for neutralization tests.

### Characterization of wild-type and chimeric SARS-CoV-2’s with variant spikes

We quantified the P1 stocks for their genomic RNA content by RT-qPCR and for their infectious titers by plaque assay on Vero E6 cells. The RT-qPCR and plaque assays were performed as previously reported.^1,23^ The ratio of viral RNA to PFU was calculated to indicate the specific infectivity of each virus.

### BTN162b2-vaccinated human sera

A panel of 20 sera was collected from 15 BTN162b2-immunized participants in the pivotal clinical trial.^6,15^ The sera were collected 2 or 4 weeks after two doses of 30 μg of BNT162b2 mRNA, spaced 3 weeks apart. As indicated in **Extended Data Table 1**, 5 of the 20 participants provided sera at both 2 and 4 weeks after the second dose of vaccine. The ages of human subjects are also presented in **Extended Data Table 1**.

### Plaque-reduction neutralization test

A PRNT_50_ was performed to measure the neutralizing titers of individual serum specimens. A detailed PRNT_50_ protocol was reported previously.^24^ Individual sera were 2-fold serially diluted in culture medium with a starting dilution of 1:40. One hundred PFUs of WT or chimeric SARS-CoV-2 with variant spike were mixed with the serially diluted sera. After incubation at 37 °C for 1 h, the serum/virus mixtures were inoculated on to 6-well plates with a monolayer of Vero E6 cells (pre-seeded the previous day). The PRNT_50_ titer was defined the minimal serum dilution that suppressed >50% of viral plaques.

### Statistical analysis

Statistical analyses were performed by Graphpad Prism 9 for all experiments as detailed in legends to individual figures.

## Data availability

Source data for generating the main figure are available in the online version of the paper. Any other information is available upon request.

## Acknowledgments

We thank the participants from Pfizer-BioNTech clinical trial C4591001, who donated the post-vaccination sera. We also thank the colleagues at Pfizer and BioNTech who developed and produced the BNT162b2 vaccine candidate. We thank Qi Yang for analysis of variant sequences. The study was supported by Pfizer and BioNTech.

## Author contributions

Conceptualization, K.U.J, U.S., X.X., K.A.S., A.M., P.R.D., P.-Y.S.; Methodology, J.L., Y.L., H.X., J.Z., S.C.W., K.A.S., H.C., A.M., K.U.J., U.S., X.X., P.R.D., P.-Y.S.; Investigation, J.L., Y.L., H.X., J.Z., S.C.W., K.A.S., H.C., M.C., D.C., K.U.J., U.S., X.X., P.R.D., P.-Y.S.; Data Curation, J.L., Y.L., M.C., D.C., X.X., P.-Y.S.; Writing-Original Draft, J.L., Y.L., X.X., P.R.D., P.-Y.S.; Writing-Review & Editing, S.C.W., K.A.S., A.M., K.U.J., U.S., X.X., P.R.D., P.-Y.S.; Supervision, K.U.J., U.S., X.X., P.R.D., P.-Y.S.; Funding Acquisition, K.U.J., U.S., P.R.D., P.-Y.S.

## Competing financial interests

X.X. and P.-Y.S. have filed a patent on the reverse genetic system of SARS-CoV-2. Y.L., H.X., J.Z., X.X., and P.-Y.S. received compensation from Pfizer to perform the project. K.A.S., H.C., M.C., D.C., K.U.J., and P.R.D. are employees of Pfizer and may hold stock options. A.M. and U.S. are employees of BioNTech and may hold stock options.

---

**Extended Data Figure 1.**
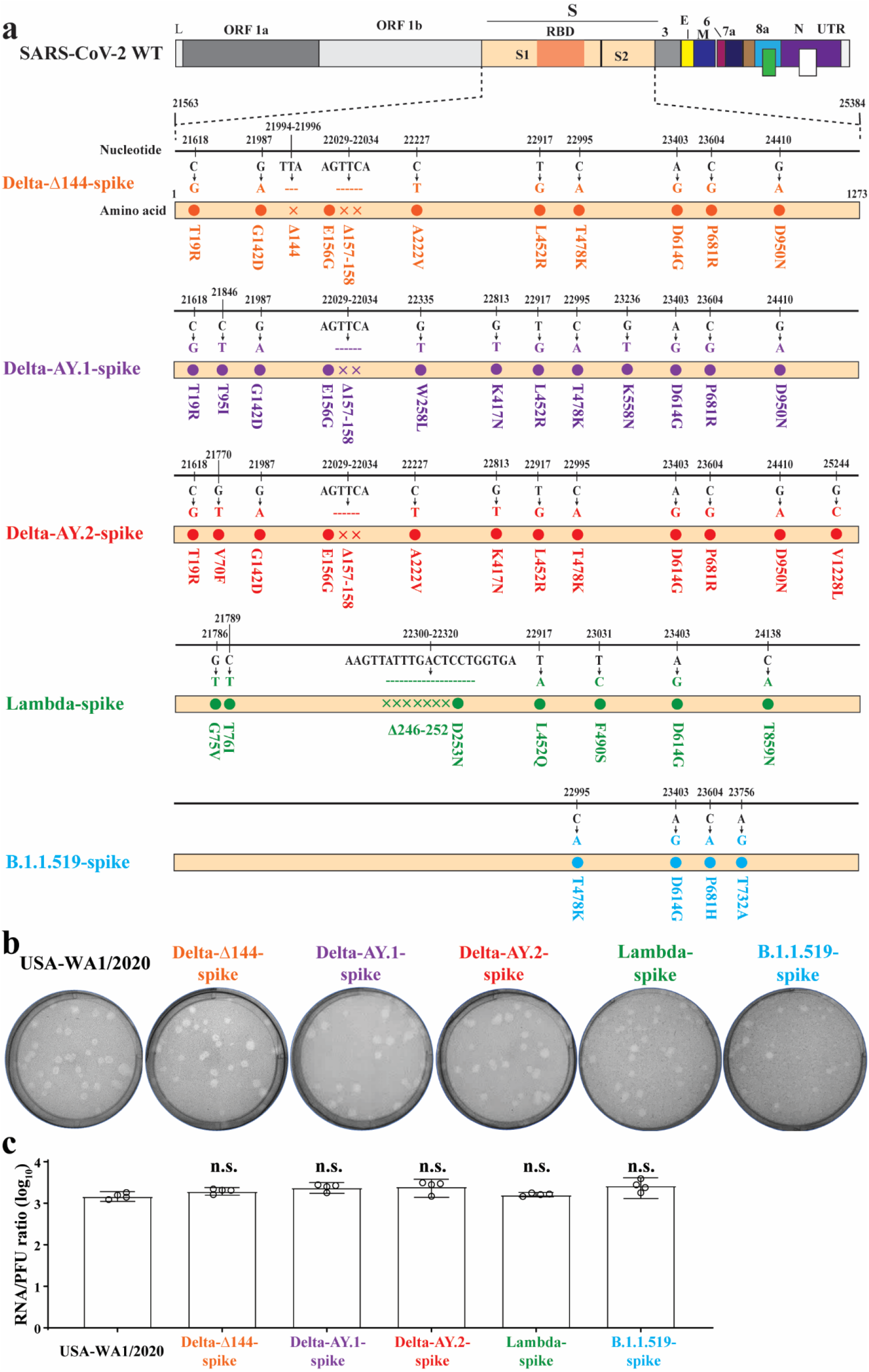
Construction and characterization of SARS-CoV-2s with variant spikes. **a,** Summary of engineered variant spike mutations. Mutations from variant spikes were engineered into USA-WA1/2020 SARS-CoV-2. Mutations and deletions are indicated by dots (•) and crosses (x), respectively. Nucleotide and amino acid positions are also shown. Regions of SARS-CoV-2 genome are indicated: L (leader sequence), ORF (open reading frame), RBD (receptor binding domain), S (spike glycoprotein), S1 (N-terminal furin cleavage fragment of S), S2 (C-terminal furin cleavage fragment of S), E (envelope protein), M (membrane protein), N (nucleoprotein), and UTR (non-translated region). **b,** Plaque morphologies of recombinant SARS-CoV-2s. Plaque assays were performed on Vero E6 cells in 6-well plates. **c,** Viral genomic RNA versus plaque-forming unit ratios (RNA/PFU) of recombinant SARS-CoV-2s. The genomic RNA content and PFU of individual virus stocks were measured by RT-qPCR and plaque assay, respectively. The RNA/PFU ratios were calculated to determine specific infectivities. Dots represent individual biological replicates from 4 aliquots of viruses (n=4, one experiment). The values in the graph represent means with 95% confidence intervals. A nonparametric two-tailed Mann-Whitney test was used to determine significant differences between USA-WA1/2020 and variant viruses. *P* values were adjusted using the Bonferroni correction to account for multiple comparisons. Differences were considered significant if *P* < 0.05; n.s. means no statistical difference.

**Extended Data Table 1.** PRNT_50_ values of twenty sera from BNT162b2-vaccinated clinical trial participants against USA-WA1/2020 and SARS-CoV-2’s with variant spikes

| *<sup>†</sup>Serum |  |  | PRNT<sub>50</sub> |  |  |  |  |  |  |  |
| --- | --- | --- | --- | --- | --- | --- | --- | --- | --- | --- |
| ID | Age | Week | <sup>‡</sup>USA-WA1/2020 |  |  | Delta-AY.1 | Delta-AY.2 | Delta-Δ144 | Lambda | B.1.1.519 |
|  |  |  | Exp1 | Exp2 | GMT |  |  |  |  |  |
| 1 | 68 | 2 | 640 | 640 | 640 | 640 | 640 | 640 | 640 | 640 |
| 2 | 67 | 2 | 160 | 80 | 113 | 80 | 80 | 160 | 80 | 160 |
| 3 | 68 | 2 | 1280 | 1280 | 905 | 640 | 1280 | 640 | 640 | 1280 |
| 4 | 65 | 2 | 320 | 320 | 453 | 320 | 320 | 320 | 640 | 640 |
| 5 | 30 | 2 | 320 | 80 | 320 | 320 | 80 | 320 | 320 | 320 |
| 6 | 23 | 2 | 640 | 320 | 453 | 320 | 320 | 160 | 640 | 640 |
| 7 | 54 | 2 | 1280 | 640 | 905 | 640 | 640 | 640 | 640 | 640 |
| 8 | 69 | 2 | 640 | 320 | 453 | 320 | 320 | 640 | 640 | 640 |
| 9 | 65 | 2 | 640 | 640 | 905 | 640 | 640 | 640 | 1280 | 1280 |
| 10 | 38 | 2 | 320 | 640 | 453 | 640 | 640 | 640 | 640 | 640 |
| 11 | 44 | 2 | 640 | 320 | 640 | 640 | 320 | 320 | 640 | 640 |
| 12 | 52 | 2 | 640 | 320 | 640 | 640 | 320 | 320 | 640 | 640 |
| 13 | 28 | 2 | 1280 | 320 | 1280 | 640 | 320 | 640 | 640 | 1280 |
| 14 | 69 | 4 | 320 | 320 | 453 | 160 | 320 | 320 | 640 | 640 |
| 15 | 68 | 4 | 320 | 320 | 320 | 640 | 320 | 640 | 640 | 640 |
| 16 | 26 | 4 | 320 | 160 | 320 | 640 | 160 | 160 | 640 | 640 |
| 17 | 54 | 4 | 640 | 640 | 640 | 640 | 640 | 640 | 1280 | 640 |
| 18 | 35 | 4 | 320 | 640 | 453 | 640 | 640 | 320 | 640 | 640 |
| 19 | 44 | 4 | 640 | 320 | 640 | 320 | 320 | 320 | 640 | 640 |
| 20 | 52 | 4 | 640 | 320 | 640 | 640 | 320 | 320 | 640 | 640 |
| <sup>§</sup>GMT |  |  | 520 | 520 | 520 | 355 | 394 | 453 | 597 | 640 |
| <sup>^</sup>95% CI |  |  | 401-674 | 393-688 | 408-663 | 258-489 | 311-500 | 346-592 | 462-771 | 519-790 |

\*Five of the twenty participants donated pairs of sera at both 2 and 4 weeks after the second dose of vaccine. The paired sera have specimen IDs of 1 and 15, 7 and 17, 8 and 14, 11 and 19, and 12 and 20.
<sup>†</sup>The serum donors were White, except for donor 10, who was Asian. All donors were of non-Hispanic/non-Latino ethnicity.
<sup>‡</sup>The data for USA-WA1/2020 are from two independent experiments. The results for other variants are from one experiment each. For each independent experiment, the individual PRNT<sub>50</sub> value is the geometric mean of duplicate plaque assay results; no differences were observed between the duplicate assays.
<sup>§</sup>GMT: Geometric mean neutralizing titers.
<sup>^</sup>95% CI: 95% confidence interval (95% CI) for the GMT.

